# Challenges in predicting protein-protein interactions of understudied viruses: Arenavirus-Human interactions

**DOI:** 10.1101/2025.04.16.649136

**Authors:** Harshita Sahni, Sarah Michelle Crotzer, Juston Moore, Steven S. Branda, Trilce Estrada, S. Gnanakaran

## Abstract

Understanding protein-protein interactions (PPIs) between viruses and human proteins is crucial for uncovering infection mechanisms and identifying potential therapeutic targets. The ability to generalize PPI predictive models across understudied viruses presents a significant challenge. In this work, we use arenavirus-human PPIs to illustrate the difficulties associated with model generalization, which are compounded by a lack of both positive and negative data. We employ a Transfer Learning approach to investigate arenavirus-human PPI by utilizing models trained on better-studied virus-human and human-human interactions. Additionally, we curate and assess four types of negative sampling datasets to evaluate their impact on model performance. Despite the overall high accuracies (93-99%) and AUPRC scores (0.8-0.9) appearing promising, further analysis indicates that these performance metrics can be misleading due to data leakage, data bias, and overfitting, especially concerning under-represented viral proteins. We reveal these gaps and assess the impact of data imbalance through standard k-fold cross-validation and Independent Blind Testing with a Balanced Dataset, leading to a drop in accuracy below 50%. We propose a viral protein-specific evaluation framework that groups viral proteins into majority and minority classes based on their representation in the dataset, allowing for comparison of model performance across these groups using balanced accuracies. This framework offers a more robust evaluation of model generalizability, addressing biases inherent in standard evaluation techniques and paving the way for more reliable PPI prediction models for understudied viruses.

## Introduction

Protein-protein interactions (PPIs) are physical contact between two or more proteins, these interactions are essential for many cellular processes. PPIs have also emerged as key elements in designing therapeutic solutions for infectious diseases [1]. High-throughput experimental techniques such as yeast two-hybrid analysis [2] and TAP-tagging [3] have produced extensive PPI datasets, fueling the application of machine learning (ML) and deep learning (DL) approaches to predict as yet unobserved PPIs. While such experimental techniques provide valuable ground-truth data, they also encounter some challenges: They are generally time-consuming and expensive. Additionally, the data they produce is potentially biased since they often struggle to capture certain types of interactions, particularly those that are weak; and in some cases, they are not able to distinguish direct interactions from multivalent associations of multiple proteins [4]. It is expected that computational approaches based on deep learning will help overcome the limitations of the experimental techniques by exposing features in PPI data that are not discernible through conventional statistical approaches, thus providing a more comprehensive and efficient solution to PPI analysis. [5–7]

Early efforts in PPI prediction employed classical ML classifiers such as support vector machines (SVM) and random forests (RF). Guo et al. [8] used SVM with auto covariance (AC) features that demonstrated effectiveness in PPI predictions in the yeast *Saccharomyces cerevisiae*. Similarly, PPI-Detect [5], ACTSVM [9], and other approaches [10] have applied SVM to binary PPI predictions. Additionally, an RF-based approach using Multi-scale Local Descriptor (MLD) features was applied to *Saccharomyces cerevisiae* and *Helicobacter pylori* data [11]. Other classifiers such as XGBoost and LightGBM have been extensively applied as well. For instance, in [12] the Chen et al. designed StackPPI that integrates XGBoost within an ensemble framework leveraging various feature sets to enhance prediction accuracy. In reference [13], the authors developed LightGBM to expedite training by removing redundant and noise features for PPI prediction across a variety of organisms, including *Caenorhabditis elegans*, *Escherichia coli*, *Homo sapiens*, and *Mus musculus*.

More recent studies on virus-host PPI predictions utilize deep learning architectures that capture complex biological sequence patterns. Architectures like recurrent neural networks (RNNs), convolutional neural networks (CNNs), and attention-based mechanisms (transformers) have shown improvements in predicting virus-human PPIs by capturing local patterns and preserving long-term dependencies in sequential data [14]. DeepViral [15], a CNN-based model, applies DL2Vec embeddings to predict interactions across various virus-human systems. In [16] Yang, et al. introduced TransPPI, a transfer learning approach that employs a Siamese CNN with a Multi-Layer Preceptron (MLP) to predict PPIs for multiple viral pathogens, including HIV, Herpes, and Dengue. Similar efforts by Chen et al. [17], and Soleymani et al. [18] utilize CNN architectures to enhance PPI prediction accuracy. Furthermore, Long short-term memory (LSTM) subnetworks [19], coupled with word2vec embeddings, have been used effectively for predicting SARS-CoV-2 PPIs with human proteins. The DeepVHPPI method [20] integrates a self-attention transformer and transfer learning to predict virus-host PPIs for Ebola, H1N1, and SARS-CoV-2. While deep learning models offer improved prediction capabilities, they are computationally intensive and require large datasets to generalize correctly [21, 22].

Positive virus-host PPI datasets typically consist of experimentally validated virus-host PPIs, which are typically sourced from public databases such as HPIDB [23], DIP [24], BioGrid [25], STRING [26], IntAct [27], ViralHostNet [28], PHISTO [29], PDB [30] and/or are extracted directly from relevant literature. A negative virus-host PPI consists of pairs that are assumed to not form a physical interaction under biological conditions. Negative virus-host PPI data, lacks a universally accepted gold standard, leading to the prevalent use of random sampling methods. In random sampling, virus-host pairs are randomly selected, assuming they do not interact if there is no evidence of interaction. However, random sampling may inadvertently introduce false negatives into the training datasets. Alternative approaches, such as Dissimilarity-Based Negative Sampling [31], have been proposed to address this issue. The Negatome database [32] has some negative data collection which is experimentally verified, but its coverage of virus-host PPI is extremely limited with few proteins from an even fewer number of viruses.

The challenge of generalizing ML models across different viruses remains a significant issue, underscoring the need of improving these predictive models to new data [14, 21, 22]. In this work, we use arenavirus-human PPIs as an example to highlight the struggles with generalization across different viruses. Arenaviruses are enveloped viruses with a segmented single-stranded ambisense RNA genome. The *Arenaviridae* family is comprised of the *Mammarenavirus* genus, members of which reside in rodent hosts and can be transmitted to humans; and three other genera (*Reptarenavirus*, *Hartmanivirus*, *Antennavirus*) whose members reside in non-mammalian hosts (e.g., snakes, fish) and cannot be transmitted to humans. *Mammarenaviruses* (hereafter referred to as “arenaviruses”) are categorized into two main geographical groups: Old World arenaviruses, prevalent in Africa and Europe, and New World arenaviruses, found in the Americas [33]. Arenaviruses cause a number of diseases, ranging from mild febrile illnesses to severe hemorrhagic fevers. These diseases are of significant public health concern, particularly in endemic regions. Some arenaviruses, such as the Lassa virus, are also considered potential bioterrorism threats due to their pathogenicity and potential for widespread impact [34, 35].

Arenavirus-human PPIs are understudied compared to viruses such as HIV, influenza, and SARS-CoV-2. The relative scarcity of experimentally generated data makes it difficult to train robust predictive models directly on arenavirus-specific data. However, transfer learning can compensate for the lack of data by using models pre-trained on other virus-human PPI. In transfer learning, a model uses the information it has learned from one task to enhance its generalization about another [36]. Transfer learning is specifically helpful in cases where data is limited. Therefore, we hypothesize that transfer learning will enable us to utilize knowledge from well-studied PPIs where more data and resources are available to improve the understanding of less-studied PPIs, like those involving arenaviruses. By training on larger and more diverse datasets, transfer learning can help models learn generalizable features relevant to arenavirus-human PPIs, potentially reducing the risk of overfitting. Moreover, this can also help reduce the need for extensive new data collection and annotation.

In this work, we explore the use of a transfer learning technique, TransPPI [16], to study and predict arenavirus-human PPIs. We leverage models that are pre-trained on data from other virus-human PPI data (e.g., from HIV, herpes, SARS-CoV-2, etc.) or human-human PPI data. With the limited availability of data for arenavirus-human PPIs, our work assesses how well models can generalize across viral families. Our study explores multiple setups, including curating various negative datasets and data balancing strategies, to understand how different testing scenarios affect model performance. We further investigate the impact of training techniques, such as k-fold cross-validation and independent testing, while addressing issues like data leakage, overfitting, and data bias. We propose a rigorous evaluation framework of virus-human PPI predictions by using a comprehensive set of evaluation metrics that includes comparing balanced accuracies for minority and majority classes and viral protein specific analysis using confusion matrix.

## Methodology

This section outlines the processes of data collection, feature representation, model training, transfer learning and evaluation metrics.

### 1. Data Collection and Preprocessing

To train machine learning models for a binary classification task, it is essential to have both positive and negative examples. Positive examples represent instances where the desired outcome is present (interacting protein pairs in a protein-protein interaction prediction task), while negative examples represent cases where the outcome is absent (non-interacting protein pairs).

#### Positive Arenavirus Dataset

To construct a robust positive PPI dataset, we aggregated arenavirus-human PPI data from several reputable databases, including HPIDB [23], DIP [24], BioGrid [25], STRING [26], IntAct [27], ViralHostNet [28], PHISTO [29] and PDB [30]. After cleaning the data by removing all the faulty and duplicate entries, our final Positive Dataset comprises 430 positive interaction pairs, representing experimentally confirmed arenavirus-human PPIs (**Table 1**). Dataset details with the protein names and uniport IDs as a csv file can be found in the supplementary material of this work.

**Table 1.** List of the sources from which the Positive Dataset was compiled and number of unique pairs found.

| Database Name | Number of pairs |
| --- | --- |
| ViralHostNet [28] | 404 |
| HPIDB [23] | 20 |
| PHISTO [29] | 3 |
| PDB [30] | 3 |

#### Negative Datasets

We utilized multiple approaches to assemble four negative datasets containing non-interacting arenavirus-human protein pairs. It should be noted that no negative experimentally verified dataset was available. Rather, these are arenavirus-human PPI pairs, for which the interactions were “unknown” or dissimilar to pairs in the positive dataset. We gathered four different negative PPI datasets. For each of the datasets we sampled a total of 4300 pairs, to keep a ratio of 1:10 positive to negative data.

We constructed four different datasets, each combining Positive Arenavirus Dataset with a different negative dataset, namely Canonical Dataset (CANON), Broad Hybrid Clustered Dataset (HYBRID-B), Refined Hybrid Clustered Dataset (HYBRID-R) and Randomized Pairing Dataset (RANDP), listed in **Table 2**.

**Table 2.**
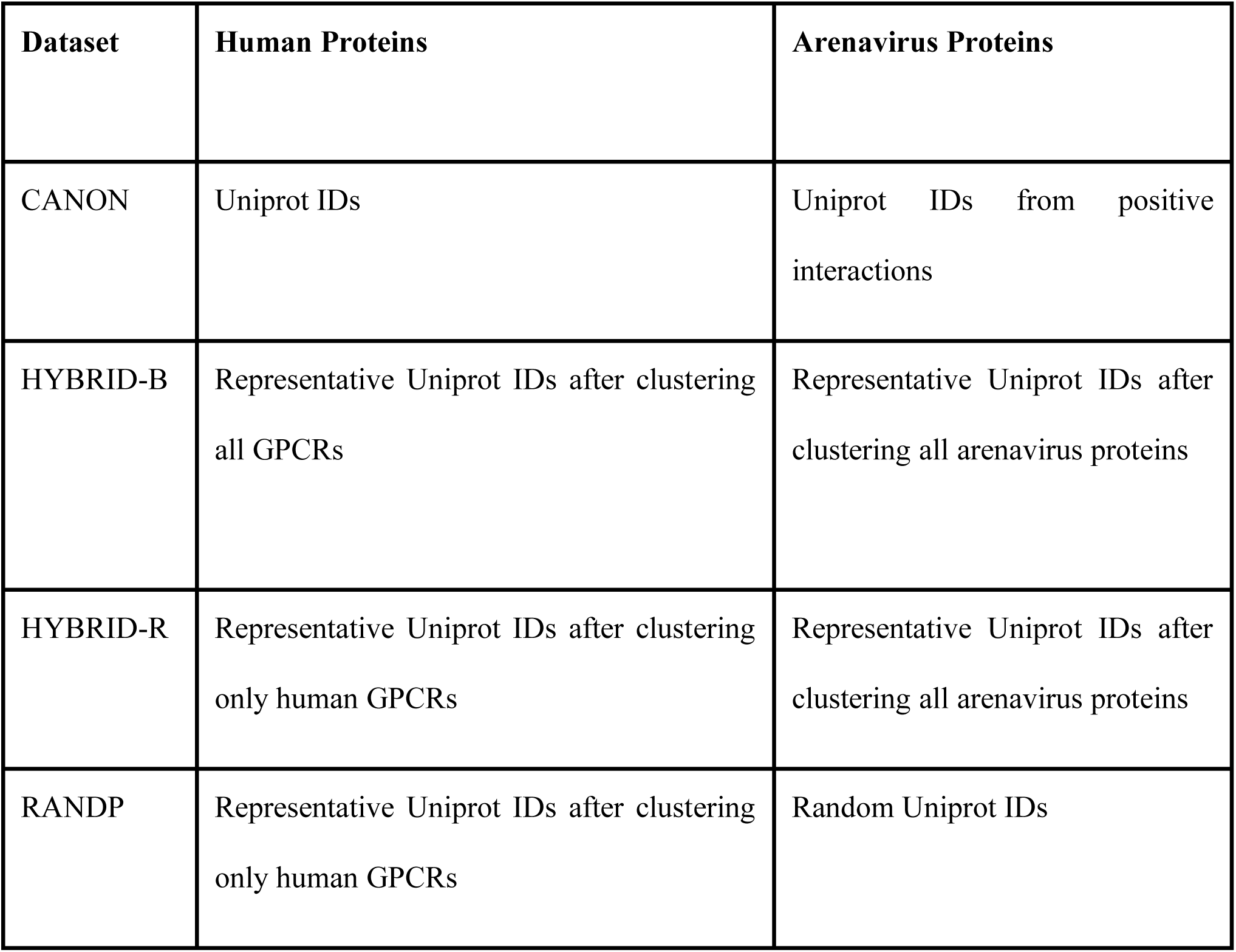
Different types of negative data sampling for arenavirus-human PPI datasets.

| Dataset | Human Proteins | Arenavirus Proteins |
| --- | --- | --- |
| CANON | Uniprot IDs | Uniprot IDs from positive interactions |
| HYBRID-B | Representative Uniprot IDs after clustering all GPCRs | Representative Uniprot IDs after clustering all arenavirus proteins |
| HYBRID-R | Representative Uniprot IDs after clustering only human GPCRs | Representative Uniprot IDs after clustering all arenavirus proteins |
| RANDP | Representative Uniprot IDs after clustering only human GPCRs | Random Uniprot IDs |

##### Negative sampling for CANON dataset

We used Dissimilarity-Based Negative Sampling [31] to collect the negative PPI pairs. This is a standard approach in the literature to sample negative pairs that are sufficiently different from positive interactions. This method operates on the principle that if a viral protein A is similar to a viral protein B by more than a threshold (30% in this work), and B interacts with human protein C (i.e., B-C represents a positive sample), then the protein pair A-C should not be classified as a negative sample. To populate this dataset, we made pairs from all arenavirus sequences present in the positive dataset and human sequences (205, 003) sourced from UniProt [37]. After performing dissimilarity-based negative sampling we kept 4300 negative pairs.

##### Negative sampling for HYBRID-B dataset

This dataset was gathered by making a biologically informed assumption. In our positive virus-human PPI datasets from other viruses including DENV, Hepatitis, SARS-CoV-2, Zika, Herpes, Influenza, Papilloma, and HIV, only 0.06% of the total pairs were identified as human GPCR interactions. Also, there were no PPIs between arenavirus protein and GPCRs in our positive dataset. Considering this limited number of GPCR’s in the positive dataset, we enriched the negative dataset with broad and diverse human GPCRs. We collected 792,466 sequences of G-protein-coupled receptors (GPCRs) and 5678 sequences of *Arenaviridae* proteins from UniProt [37].

To ensure data quality and minimize information leakage, we applied clustering algorithm cd-hit [38] to cluster similar sequences with specific similarity thresholds. For the GPCRs, we used a 40 percent sequence similarity threshold, grouping sequences that share 40 percent or more similarity into a single cluster. For arenaviruses, we applied a sequence similarity threshold of 70 percent. To create this dataset, we paired a representative sequence from each GPCR cluster with one from each arenavirus protein cluster. This step effectively removed duplicate or highly similar sequences, ensuring a diverse and representative dataset for model training and evaluation. It is termed as broad and hybrid as the cd-hit clustering is applied to broad category of GPCR proteins pairing with arenavirus proteins.

##### Negative Sampling for HYBRID-R

We further refined the 792,466 sequences of GPCRs by filtering and selecting only the human GPCRs (yielding a subset of 2869 sequences) and then applying cd-hit clustering with a 40 percent sequence similarity threshold. This additional dataset focuses on human GPCRs, allowing for a more targeted comparison. To create this dataset, we paired the representative sequences from human GPCR and arenavirus protein clusters, as with the HYBRID-B set. Since, it is built upon hybrid clustering approach but applies an additional refinement in selecting only human-GPCR proteins, we name it HYBRID-R.

##### Negative Sampling for RANDP

For a representative set of human protein sequences, we used the same set of sequences as in HYBRID-R. However, for a diverse set of arenavirus protein sequences, instead of clustering we randomly sampled sequences from the complete *Arenaviridae* dataset. We then paired each representative human-GPCR sequence with a randomly selected arenavirus protein sequence. This dataset is named RANDP as we randomly select arenavirus proteins and paired them with human-GPCRs.

### 2. Feature Representation

We computed Position-Specific Scoring Matrices (PSSMs) for each protein sequence using PSI-BLAST [39] for feature extraction. PSSMs provide a detailed representation of sequence conservation and evolutionary information by highlighting amino acid substitutions and their frequencies [40]. These metrics serve as informative input features for the PPI prediction model, allowing it to learn complex sequence patterns associated with protein interactions. We used PSI-BLAST to compute PSSM with an e-value of 0.001 and three iterations. The computation of PSSM profiles was distributed across multiple processing units using the Message Passing Interface (MPI) framework [41]. Using MPI, the workload is distributed across multiple nodes, each handling a subset of the sequences. We scaled our pipeline on about ∼79,000 sequences up to 512 processes across four nodes. This distributed approach facilitated efficient processing of the large dataset, reducing computation time and resource requirements. By distributing the computations across multiple nodes and multiple CPU units we made better use of the computational infrastructure and achieved a significant speedup.

### 3. Model Training with TransPPI

In this study, we adopt the TransPPI method [16], which involves initially training on diverse virus-human PPI datasets from a range of viruses including Dengue, Hepatitis, HIV, SARS-CoV-2, Zika, Herpes, Influenza, and Papilloma viruses and subsequently evaluating the model’s ability to predict PPIs across these viral systems. In TransPPI, positive PPI examples are curated from experimentally validated databases such as HPIDB, while negative samples are generated by dissimilarity based negative sampling, using a 1:10 positive-to-negative ratio. The model architecture consists of four convolutional layers, which are well-suited for capturing hierarchical patterns in protein sequences. This initial training phase aims to establish a foundational understanding of general interaction patterns, enabling the model to learn from a broad range of biological contexts. TransPPI has demonstrated improved cross-viral generalization and higher predictive performance compared to traditional methods. TransPPI achieved AUPRC scores ranging from 0.329 (SARS-CoV-2) to 0.974 (HIV) across different virus-human PPI datasets.

### 4. Transfer Learning and FineTuning

Transfer learning was employed to adapt the pre-trained TransPPI model to the arenavirus PPI predictions, taking advantage of its ability to generalize knowledge from other PPI datasets. Two key transfer learning approaches were utilized:

#### Fine Tuning

In this approach, the pre-trained model’s weights (trained on virus-human protein pairs (e.g. HIV-human etc.)) were initialized as a starting point, and all layers of the model were subsequently finetuned using the arenavirus-human PPI datasets as listed in **Table 2**. This approach allows the model to adjust all layers and learn features specific to the new dataset, improving its predictive performance on the target interactions.

#### Frozen Layers

In this approach, a subset of the pre-trained model’s layers, particularly the initial convolutional layers, were frozen, meaning their weights were not updated during training on the arenavirus-human PPI datasets. This technique preserves the generalized features learned from the broader dataset, while only the final layers of the model are finetuned to capture dataset-specific nuances. Freezing layers helps retain previously acquired knowledge, preventing overfitting to the limited data of the target system.

### 5. Evaluation Metrics

We evaluated the performance of the models on arenavirus-human PPI datasets listed in **Table 2**. using a set of metrics that provide a complete understanding of the model’s predictive capabilities. The evaluation metrics included accuracy (ACC), which measures the percentage of interactions that were classified correctly, but does not provide insight on unbalanced data; area under the precision-recall curve (AUPRC) measures how well the model distinguish between interacting and non-interacting pairs; precision quantifies how many interacting pairs are actually correct; and recall indicates the proportion of true interactions that were predicted correctly. Matthews correlation coefficient (MCC) measures the correlation between the predicted and actual classifications. The Balanced Accuracy is calculated by averaging recall across various classes. TP stands for True Positive (number of correctly predicted interacting pairs), FP: False Positive (number of incorrectly predicted interacting pairs), TN: True Negative (number of correctly predicted non-interacting pairs), FN: False Negative (number of incorrectly predicted non-interacting pairs)

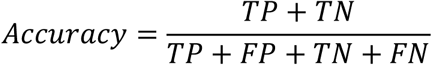

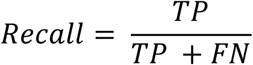

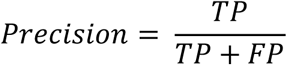

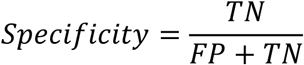

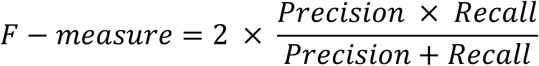

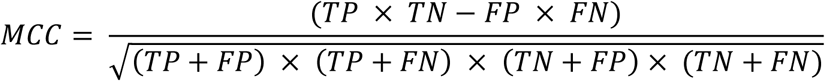

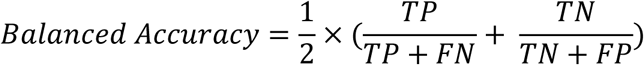

#### Viral protein specific evaluation

In addition to traditional metrics, we introduce viral protein specific evaluation. This approach focuses on determining the model’s ability to predict interactions specific to each viral protein category and making comparisons to others. We further divided each viral category (sequence) into majority and minority classes based on the total count of associated positive interactions. Viral proteins with number of samples above the median count were assigned to the majority class, while those with fewer were designated as minority class. We calculated the balanced accuracy for both the majority and minority viral protein classes to see how well the model performed on underrepresented viral proteins (minority class) and well-represented viral proteins (majority class). We also computed MCC and overall balanced accuracy to ensure fair assessment in case of imbalanced and skewed data.

## Results

### 1. Influence of negative dataset

First, we investigated the effect of the negative datasets on model performance, considering that we had only a limited number of interactions in the positive dataset. We aim to understand how the model distinguishes between the fixed set of positive samples and each specific type of negative data. This enhances our understanding of the model’s behavior, robustness, and sensitivity to different types of negative pairs. We curated four distinct arenavirus-human datasets (CANON, HYBRID-B, HYBRID-R, RANDP), with different negative sampling strategies, as described in Sub-section #1 in Methodology and **Table 2**.

We initially trained eight ML models on virus-human PPI datasets, specifically: Human-Dengue (DENV), Human-Hepatitis, Human-HIV, Human-SARS-CoV-2 (SARS2), Human-Zika (ZIKV), Human-Herpes, Human-Influenza, and Human-Papilloma and later finetuned with our four curated arenavirus-human datasets, as discussed in Sub-section #1 in Methodology, using 5-fold cross-validation (CV). The benchmark of their performances in terms of AUPRC is provided in **Table 3**. Regardless of the virus-human dataset used for initial training, or the specific arenavirus dataset used for finetuning, the AUPRC remained stable; around 0.8 for the CANON dataset and approximately 0.9 for the other three (Hybrid-B, Hybrid-R, and RANDP) datasets. Overall, the results were consistent mainly across all three datasets. This level of stability, despite variations in training data and negative sampling strategy, may reflect an over-optimistic evaluation. Slightly lower AUPRC (0.8) values for CANON Dataset can be explained by the fact that the negative samples are generated using Dissimilarity-based negative sampling. This introduces more challenging negatives that are closer in similarity to the positive dataset, since the same virus can have both interacting and non-interacting pairs with different human proteins. In the other datasets, the negative samples are generated under a biologically informed assumption that positive virus-host PPI datasets had minimal interactions with human GPCRs. Thus, the positive and negative samples come from disjoint sets, likely making it easier for the model to classify correctly.

**Table 3.** AUPRC for models initially trained on various source systems [Human-Dengue (DENV), Human-Hepatitis, Human-HIV, Human-SARS-CoV-2 (SARS2), Human-Zika (ZIKV), Human-Herpes, Human-Influenza, Human-Papilloma] and finetuned with different arenavirus -human PPI dataset as detailed in Table 2.

| **Source** | **CANON**<br>**Dataset** | **HYBRID-B**<br>**Dataset** | **HYBRID-R**<br>**Dataset** | **RANDP Dataset** |
| --- | --- | --- | --- | --- |
| Human-DENV | 0.847 | 0.995 | 0.993 | 0.998 |
| Human-Hepatitis | 0.851 | 0.994 | 0.994 | 0.997 |
| Human-HIV | 0.847 | 0.996 | 0.997 | 0.997 |
| Human-SARS2 | 0.835 | 0.995 | 0.996 | 0.998 |
| Human-ZIKV | 0.857 | 0.994 | 0.993 | 0.998 |
| Human-Herpes | 0.816 | 0.996 | 0.993 | 0.998 |
| Human-Influenza | 0.848 | 0.995 | 0.995 | 0.999 |
| Human-Papilloma | 0.84 | 0.995 | 0.995 | 0.998 |

### 2. Transfer Learning from other virus-human PPI interactions on Canonical Dataset

After discovering that the Canonical Dataset posed greater complexity and lowest AUPRC, we selected it to further investigate the effectiveness of transfer learning capability from various systems consisting of virus-human PPI datasets. These include Human-Dengue (DENV), Human-Hepatitis, Human-HIV, Human-SARS-CoV-2 (SARS2), Human-Zika (ZIKV), Human-Herpes, Human-Influenza, and Human-Papilloma. Initially, the models were trained separately on each of the above-mentioned datasets using 5-fold cross-validation (CV). The performance of Canonical Dataset consisting of arenavirus-human PPI pairs was evaluated in detail under two distinct conditions: finetune and frozen, as shown in **Figure 1 and S1**. In finetuning, all layers of the CNN and MLP were optimized to adapt to arenavirus data. Conversely, in the frozen setting, only the MLP layers were retrained using arenavirus data. The bars in **Figure 1** illustrate the performance measurements, including AUPRC (Area Under Precision-Recall Curve), accuracy, precision, and recall for each of the finetuned models. Notably, all the models achieve relatively consistent accuracy across these viruses, indicating stable performance on these datasets. The accuracy for finetuned setting ranged from 95.2% to 96.2%, while for frozen setting it ranged from 93.5% to 95.5%.

**Figure 1:**
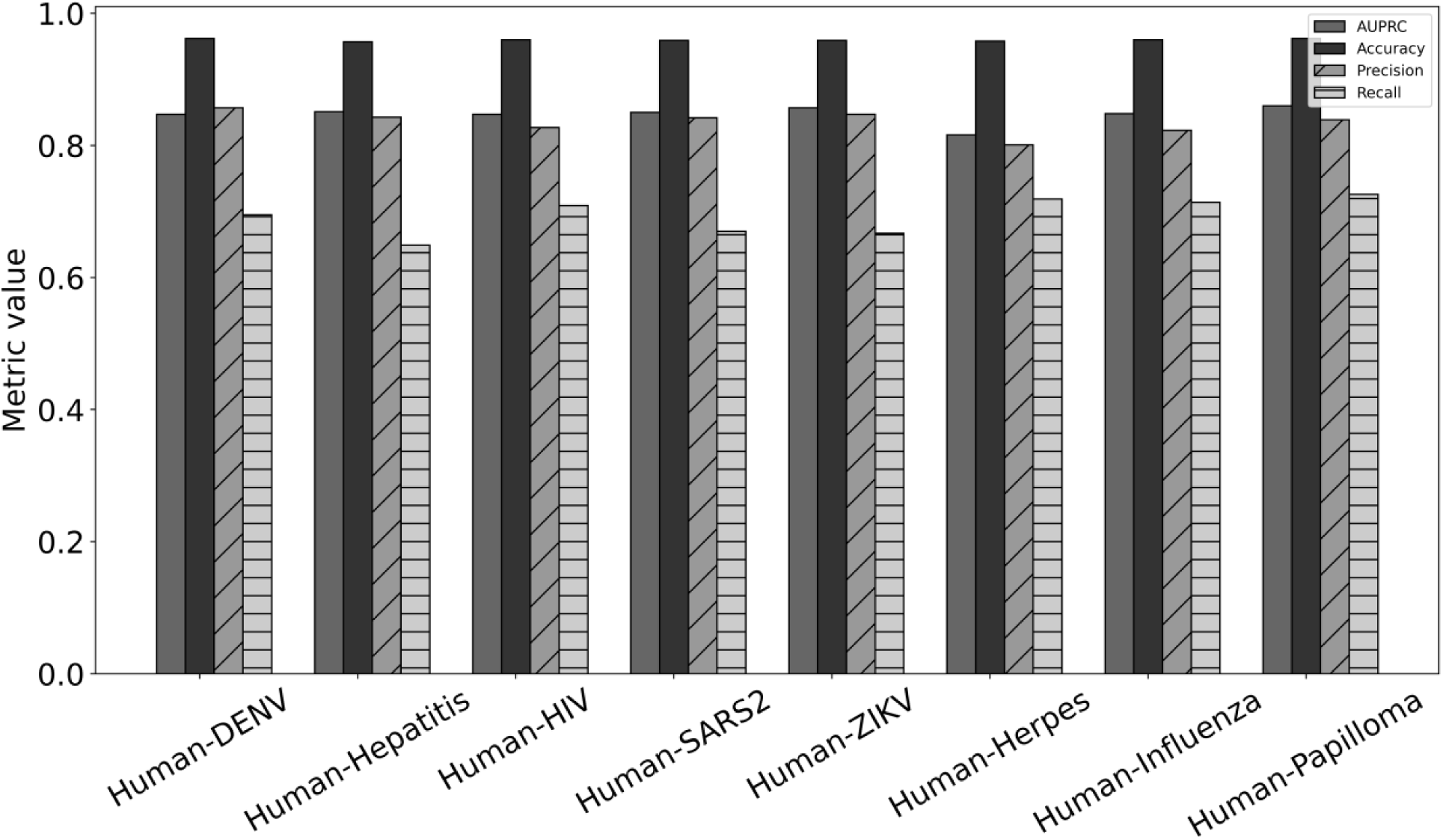
AUPRC, Accuracy, Precision and Recall for models initially trained on various source systems [Human-Dengue (DENV), Human-Hepatitis, Human-HIV, Human-SARS-CoV-2 (SARS2), Human-Zika (ZIKV), Human-Herpes, Human-Influenza, Human-Papilloma] and finetuned on Canonical Dataset.

Despite variations in the number of experimentally verified PPIs across various systems, with HIV having the highest at 9880 and SARS-Cov-2 the lowest at 568, all systems have AUPRC values that fell within the 0.8 range. The precision values across all systems are consistently high ranging from 0.801 to 0.857. However, in all scenarios, we observe high precision and low recall (0.649 - 0.726), indicating that the model is precise but has missed many actual positive instances, which are the interacting pairs. We caution that high scores alone should not be interpreted as conclusive evidence of predictive power. Since the model pretrained on HIV-human dataset contains the highest number of PPI pairs and produces one of the highest accuracies (96%), we selected this model as representative for further evaluations.

### 3. Transfer Learning from other human-human PPI interactions

Using virus-human PPI datasets initially gave us high performance, prompting us to further investigate whether using human-human PPI data could make a difference or similarly identify arenavirus-human PPI interactions. Next, we used human-human PPI data to support the prediction of arenavirus-human PPIs, leveraging the availability of vastly more experimentally verified data. We initially trained the model on the human-human PPIs included in the PAN human dataset [42] and then finetuned the model on the arenavirus-human PPI datasets listed in **Table 2**. The PAN human dataset captures a wide diversity of protein sequences, serves as a gold standard dataset for use in the analysis of human proteins, and was first used [42] for the prediction of human-human PPIs. There are 3899 pairs out of 2502 proteins in a positive dataset; less than 25% of all proteins have the similar protein sequence. The positive dataset was compiled from the reference database, HPRD (2007 edition) [43]. The negative dataset consists of 4,262 protein-protein pairs derived from a total of 661 proteins, with each protein having less than 25% identity with the others.

The model’s performance across various types of arenavirus-human PPI datasets (CANON, HYBRID-B, HYBRID-R, RANDP) is illustrated in **Figure 2**, with each type depicting similar outcomes when the model was initially trained on a human-human dataset, as compared to using other viral-human datasets **(Table 3)**. A slight increase in precision and AUPRC was observed for CANON Dataset. Specifically, the AUPRC improved from 0.847 to 0.852, and precision rose from 0.827 to 0.858 when the model was initially trained on HIV and human-human datasets, respectively, and then finetuned on the CANON Dataset. In summary, the model pre-trained on human-human PPIs achieved performance comparable to those pre-trained on HIV and other viral datasets. This indicates that regardless of the pre-training dataset and finetuning set, the model consistently produced high performance (AUPRC ∼0.8) for CANON dataset and AUPRC ∼0.9 for the other three arenavirus-human datasets.

**Figure 2:**
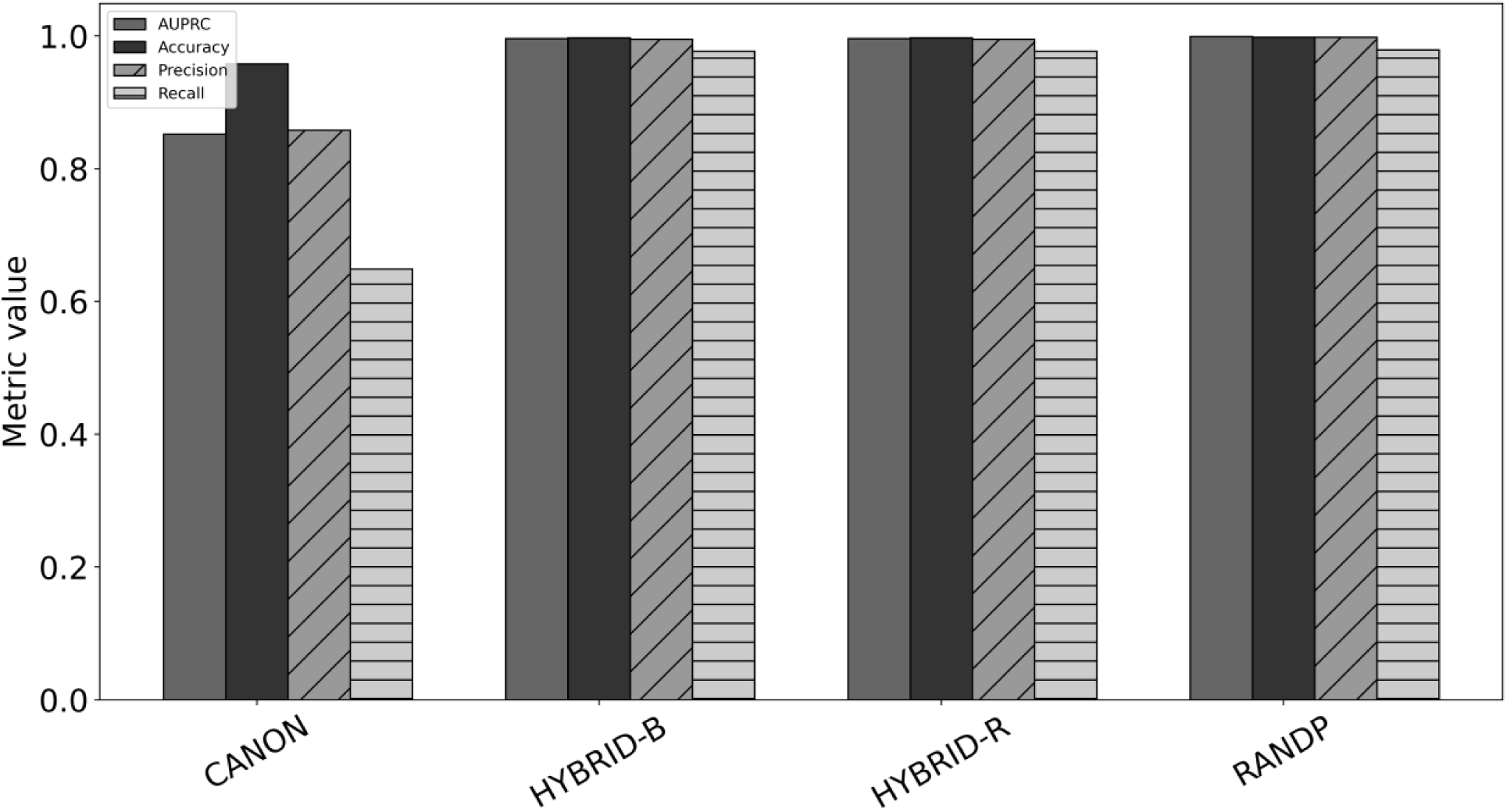
AUPRC, Accuracy, Precision and Recall for models initially trained on a human-human PPI dataset and finetuned on four arenavirus-human PPI Datasets 1-4, listed in **Table 2**.

However, high performance as measured by accuracy and AUPRC does not imply models true generalizability. The above-discussed results seem encouraging on the surface, but they can be misleading due to factors like data bias and model overfitting. Therefore, we critically examine these factors to better understand the true effectiveness and limitations of our model.

### 4. Bias from the Positive arenavirus-human PPI dataset

The Positive Dataset encompasses four categories of arenavirus proteins: Polymerase L (L), Nucleoprotein (NP), Glycoprotein (GP), and Ring Finger Protein Z (Z). As illustrated in **Figure 3**, the distribution across these categories is highly imbalanced. Specifically, the L category includes 227 samples, in each case featuring the same viral protein sequence. Similarly, the NP category includes 188 samples, in each case featuring the same viral protein sequence (both from LCMV Armstrong). Together, the L and NP samples account for 96.5% of the protein pairs comprising the Positive Dataset. In contrast, the GP category includes six samples with three different arenavirus protein sequences; and the Z protein category includes nine samples with six different arenavirus protein sequences. Therefore, the Positive Dataset includes only 11 unique arenavirus protein sequences in total **(Table 4).**

**Figure 3:**
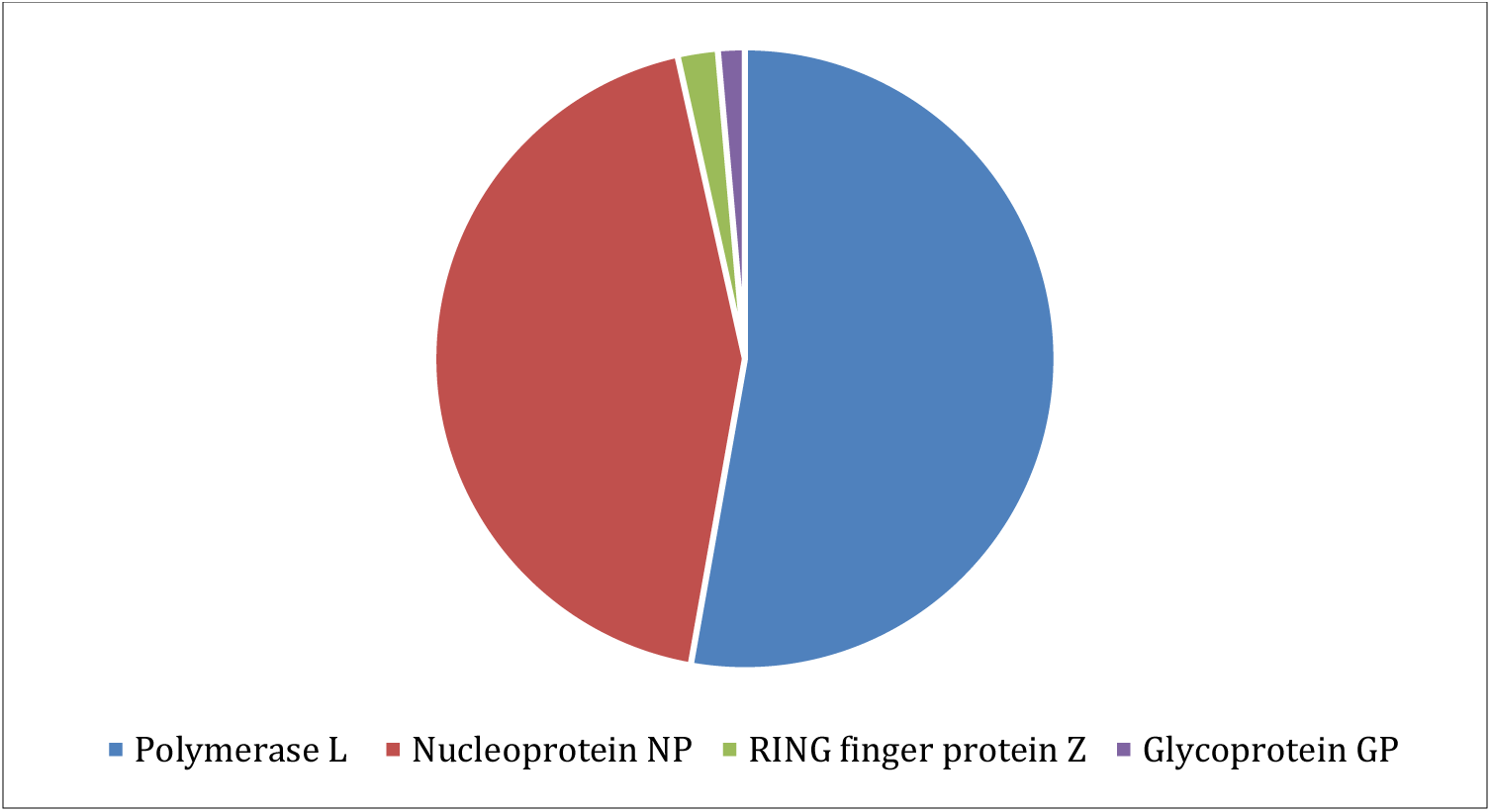
Distribution of viral proteins in the Positive Dataset, showing the number of arenavirus-human protein pairs that include the viral polymerase L, nucleoprotein NP, glycoprotein GP, and RING finger protein Z.

**Table 4.** Distribution of viral proteins in the Positive Dataset, depicting the total number of arenavirus-human PPIs for L, NP, GP, and Z, and their respective unique viral protein counts.

| Name of Type | Number of interacting pairs | Unique viral protein sequences |
| --- | --- | --- |
| Polymerase L (L) | 227 | 1 |
| Nucleoprotein (NP) | 188 | 1 |
| Glycoprotein (GP) | 6 | 3 |
| RING finger protein Z (Z) | 9 | 6 |

This imbalance in the Positive Dataset poses several challenges. The overrepresentation of certain viral proteins (NP and L; majority classes) may lead to biased models that struggle to generalize effectively to underrepresented viral proteins (GP and Z; minority classes) as discussed in Sub-section #5 in Methodology. When trained on such skewed data distributions, machine learning algorithms are susceptible to overfitting to the majority classes. Furthermore, the limited diversity within categories especially in the cases of the L and NP, each of which contains only one unique arenavirus protein sequence, restricts the model’s ability to learn a comprehensive range of interaction patterns.

Evaluating models on imbalanced datasets using metrics such as accuracy and the AUPRC can be misleading, as these metrics may not accurately reflect performance across all viral proteins (categories). The results discussed above demonstrate high performance; however, such performance can be deceptive, as the model may be overfitting to the majority class. Additionally, employing k-fold cross-validation in these contexts may fail to preserve the class distribution in each fold. Consequently, the training and testing sets may not adequately represent the overall imbalance of the dataset. As a result, models may perform well in the majority class but poorly in minority classes. This was demonstrated by making predictions on a completely independent arenavirus-human PPI dataset gathered from indirect experimental associations **(see Supplement Table S1),** which was not included in the training set. This dataset contains a diverse set of arenavirus proteins from majority and minority classes. Note that none of the pairs were in the Positive Dataset. Only one human protein and three viral proteins had interactions with a different human protein in the Positive Dataset. The model was initially trained on HIV-human PPI and finetuned on CANON dataset. Since our CANON dataset contains more interacting samples involving NP, the model tends to predict samples from this testing set as ‘interacting’ pairs whenever NP is present (five out of six times). In contrast, all nine pairs containing Z in this test dataset are predicted as non-interacting since it was underrepresented in the original Positive Dataset. This clearly indicates data bias and overfitting imposed by the majority and minority distributions in the CANON dataset, **Table S1**.

#### Viral protein specific evaluation

To better analyze the challenges associated with the imbalance in the Positive Dataset, we developed a viral protein specific evaluation, as detailed in Sub-section #5 in Methodology. Our approach treats each viral protein sequence as a distinct category. Based on the total number of samples in the positive dataset, we categorized the dataset into majority (NP and L) and minority (GP and Z) classes.

The overall accuracy metric can be misleading in imbalanced settings. Therefore, to provide a comprehensive assessment of the model’s performance we compared the overall accuracy with the balanced accuracy for the entire dataset. Then, to ensure a fair comparison across these imbalanced classes, we calculated the balanced accuracy for both the majority and minority classes within the CANON dataset, as presented in **Table 5**. To further dissect performance at the individual protein level, we analyzed the confusion matrices for each viral protein category, as illustrated in **Figure S2**. This granular analysis highlights whether any specific viral protein category is neglected or overshadowed by more dominant ones. We adopt this comprehensive evaluation strategy to detect and mitigate potential biases, ensuring that minority class viral proteins are appropriately represented in our predictive modeling.

**Table 5.** Overall and Balanced accuracies for CANON dataset under Standard k-fold, Independent Blind Testing and Independent Blind Testing with Balanced Dataset.

|  | <u>Overall</u><br><u>Accuracy</u> | <u>Balanced</u><br><u>Accuracy</u> | <u>Overall</u><br><u>Majority</u><br><u>Accuracy</u> | <u>Overall</u><br><u>Minority</u><br><u>Accuracy</u> | <u>Balanced</u><br><u>Majority</u><br><u>Accuracy</u> | <u>Balanced</u><br><u>Minority</u><br><u>Accuracy</u> | <u>MCC</u> |
| --- | --- | --- | --- | --- | --- | --- | --- |
| Standard k-fold | 0.9600 | 0.8472 | 0.8481 | 0.9969 | 0.8198 | 0.6665 | 0.7443 |
| Independent Blind Testing(1:10) | 0.9076 | 0.5138 | 0.6510 | 0.9921 | 0.5078 | 0.7305 | 0.1019 |
| Independent Blind Testing with Balanced Dataset(1:1) | 0.4884 | 0.4884 | 0.1602 | 0.9292 | 0.5012 | 0.7397 | -0.065 |

After training the initial model on the HIV-human PPI dataset, we finetuned and evaluated the model on the CANON arenavirus-human PPI dataset in three different ways and analyzed the results using viral specific evaluation framework and balanced accuracies as explained below:

1. <u>Standard k-fold Analysis</u>: We utilized the model that was initially trained on the HIV-human dataset and later finetuned on CANON dataset. The finetuning was conducted using 5-fold cross-validation (i.e. we finetuned five independent models). In this case, we maintained a 1:10 ratio of positive to negative samples, resulting in a total of 430 positives and 4,300 negative arenavirus-human pairs. The distribution of samples in negative dataset was balanced across each of the 11 arenavirus proteins, ensuring that each virus had a similar number of negative samples, ranging between 390 and 401 per arenavirus sequence. To conduct viral specific analysis, we analyzed and compared each of the 11-arenavirus protein sequences separately to one another in terms of True Positive (TP), True Negative (TN), False Positive (FP) and False Negative (FN) and balanced accuracies. The model was fairly accurate in predicting both interactions and non-interactions for L and NP, the arenavirus proteins present in the majority class samples. The TP count was 164 out of 227 (72.2%) for L, and 136 out of 188 (72.3%) for NP, as depicted in the orange bars in **Figure 4(a)**. However, performance declined for the minority classes, only 5 out of 9 (55.5%) interactions were correctly predicted for Z, and none were identified for GP (0 out of 6), as shown by the blue bars in **Figure 4(a)**. The accuracy under this setting is 96% but the balanced accuracy dropped to 84.72%. The combined accuracy of the majority classes (L and NP) is 84.81% as illustrated in **Table 5**. This further suggests that the model performance is driven largely by the majority class.
2. <u>Independent Blind Testing</u>: We utilized the models that were initially trained on the HIV dataset and then finetuned them on data from nine different viruses, reserving all data from the remaining two viruses exclusively for testing. As with the standard k-fold approach, independent blind testing was conducted with 5-fold cross-validation and a 1:10 negative-to-positive sample ratio. Importantly, for testing the two majority classes, we included data for one viral protein in the training set and the other in the testing set. This independent testing framework served as a blind test to evaluate the model’s generalizability to unseen viral proteins. In the case for L, the model’s TP rate dropped to 3.72% as compared to 72.3% in standard k-fold setting. The model was able to predict all of the non-interacting pairs correctly but, none of the interacting pairs were predicted correctly, as depicted with orange bars in **Figure 4(c,d)**. This indicates that the model struggled with recognizing actual interactions in an independent testing scenario. For the minority class Z of arenavirus proteins, only 7 out of 15 interactions (46.7%) were correctly predicted. For minority class GP, there were still no interactions that were predicted correctly, as shown in blue bars in **Figure 4(c)**. In this case, the model accuracy dropped to 90.76% and yielded a balanced accuracy of 51.38%, reflecting a decline in overall performance compared to standard k-fold testing. The majority class accuracy is 65.10%, but it is all driven by the correct predictions from the non-interaction pairs, as presented in **Table 5**.
3. <u>Independent Blind Testing with Balanced Dataset:</u> We repeated the independent testing using a dataset with a 1:1 negative-to-positive sample ratio. For this case, we maintained a 1:1 ratio of positive to negative samples, resulting in a total of 430 arenavirus-human interaction pairs. Each virus category in the negative dataset contributed approximately 33 to 34 pairs, ensuring a balanced representation across all viruses. This balanced dataset allowed us to examine how the models perform when the interacting and non-interacting distribution is equal, or if the model can still learn from less data. In this case, the predictive power for interactions of minority classes increased, as 8 out of 15 interactions (53.3%) were classified correctly. For GP, 1 out of 6 interactions are predicted correctly. However, no interacting pairs were predicted correctly (TP) in the case of L, and only one interacting pair was classified correctly in the case of NP, as depicted by orange bars in **Figure 4(e)**. We note that the five viral protein sequences in minority that are included in correctly predicted PPIs were either all paired with the same human protein sequence or are highly similar to viral protein sequences in the training set. The overall accuracy drops sharply to 48.84%, suggesting that Independent Blind Testing with Balanced Dataset has a substantial impact on overall performance. The balanced majority class accuracy (50.12%) and balanced minority class accuracy (73.97%) further emphasize the model’s difficulty in predicting interactions for positive class **(Table 5)**.

**Figure 4** summarizes our finding and **Figure S2** presents viral specific confusion matrix for all 11 arenaviruses. Overall, these results illustrate the model’s strength in identifying non-interactions across different cases (TN) but reveal challenges in identifying interacting pairs or maintaining high TP rates, especially in independent tests with balanced or imbalanced data. Also, **Table S2** presents the total count of positive and negative samples involving each of the arenavirus protein in 1:10 and 1:1 positive to negative ratios.

**Figure 4.**
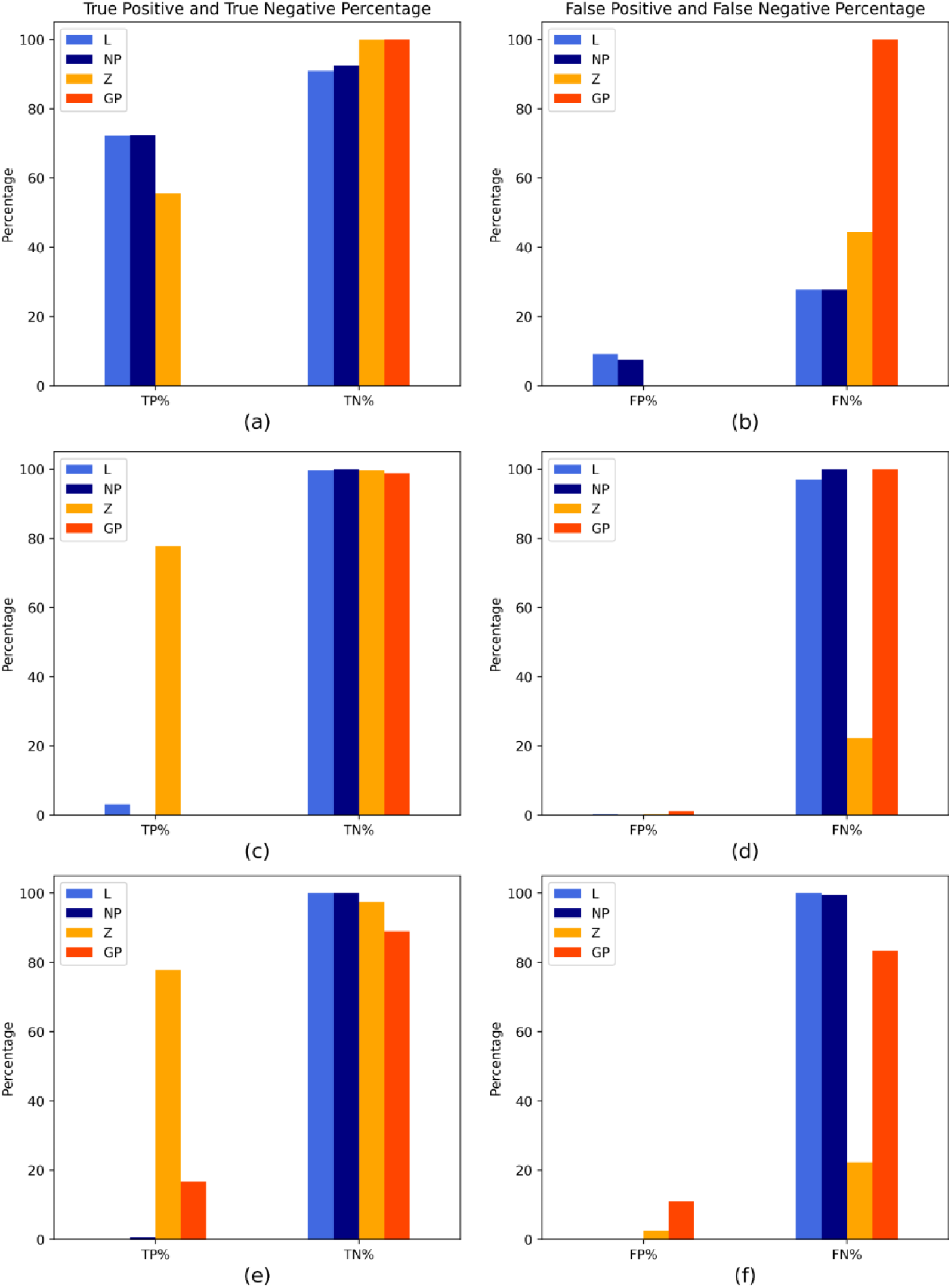
(a,c,e) shows True Positive (TP) and True Negative (TN) percent for each viral protein category, higher the better. Sub-figures (b,d,f) shows False Negative (FN) and False Positive (FP) percent for each viral protein category, lower the better Note: (a,b) represents Standard k-fold testing (c,d) Independent Blind Testing (e,f) Independent Balanced Blind Testing.

## Discussion

In this study, we employed transfer learning to address the challenge of limited experimental data in predicting virus-human protein-protein interactions (PPIs) for arenaviruses. We used host protein interactions with well-studied viral systems (HIV, DENV, SARS2, Hepatitis, etc.), as well as human-human PPIs, as the foundational data. We further evaluated four datasets with distinct negative sampling strategies, allowing us to examine their behavior in different testing scenarios. These negative datasets were based on either dissimilarity-based sampling (CANON) or biologically informed non-interacting pairs (HYBRID-B, HYBRID-R, RANDP), as provided in **Table 2**.

In our analysis of different negative datasets, the best-performing model was the one finetuned on the RANDP dataset, which shared the same positive samples as the CANON dataset but used randomly paired arenavirus proteins for the negative dataset. Compared to the model finetuned on the HIV-human dataset, the RANDP-finetuned model showed superior generalization to arenavirus-specific interactions. Virus-specific evaluation further highlighted its advantages over the CANON-finetuned model: for the majority classes, the RANDP model achieved 100% accuracy for class L (227/227) and 99% for class NP, significantly outperforming the CANON model, which reached only 72.2% for L and 72.3% for NP. For the minority classes, RANDP correctly identified 66.6% of class Z interactions, while the CANON model achieved just 55.5%; both models failed to predict any interactions for class GP, **Table S3**. We further evaluated RANDP-finetuned model on PPI pairs derived from indirect experimental associations. In this setup, the model correctly predicted 4 out of 6 interactions from the majority class (NP) and also identified 4 out of 9 interactions from the minority class (Z) as compared to model finetuned on CANON dataset that predicted 5 out of 6 interactions from the majority class (NP), and none from the minority class (Z), **Table S1.** These results indicate that the RANDP strategy not only reduces overfitting seen in CANON but also enables better generalization across both major and minor viral protein classes.

We observed that all models, regardless of the initial training dataset or the negative sampling strategy employed, demonstrated comparable performance (**Figure-1, Table-3**). However, the consistently high accuracy and AUPRC metrics across experiments do not reflect the true generalizability or robustness of the models. To address this, we adopted a viral-specific evaluation framework that divides each viral sequence into majority or minority classes based on the data distribution and focused on balanced accuracy to provide a more nuanced assessment of model performance. We further compare each viral category using a confusion matrix or correct/incorrect interacting and non-interacting pairs predicted by the model (TP, FP, TN, FN).

Our findings highlight the need for greater granularity in predicting protein-protein interaction, especially when using dissimilarity based sampling or random sampling techniques for negative data with an imbalanced data ratio. We strongly recommend that future research adopt evaluation strategies that go beyond aggregate metrics and follow a set of guidelines to improve the reliability and generalizability of PPI prediction models. First, the adoption of a viral-specific evaluation framework is essential. Models should either assess each viral protein individually or with further division into majority and minority classes based on the data distribution as discussed in the Sub-section #5 in Methodology, to understand performance across differently represented protein categories better. Second, we emphasize the importance of reporting balanced accuracy and MCC, especially when working with imbalanced datasets. Furthermore, we also recommend using loss functions like focal loss [44] to penalize errors in minority classes more heavily and help address class imbalance during training. Together, these strategies will contribute to building more balanced and confident models, ultimately improving the robustness of PPI predictions.

Our study, similar to some other deep learning-based predictions of PPIs [11, 16, 45–48], demonstrates remarkable prediction accuracies, generally ranging from 95% to 99%. However, these impressive results are often achieved on data that are randomly split into training and test sets through k-fold cross-validation. This practice inadvertently inflates performance metrics due to training data leakage. Park and Marcotte [49] also showed that random splitting can result in the same proteins appearing in both the training and test sets, which compromises the independence of these sets. Consequently, the model learns specific patterns from the training data that directly apply to the test data, leading to overestimating the models true predictive power [46].

Similar concerns arise when predicting arenavirus-human PPIs. The high accuracy of 93-99% observed in this study might be influenced by several factors that parallel those identified in the broader deep learning literature [45, 46] [[50]. The limited diversity of the arenavirus-human PPI dataset (Positive Dataset), which contains only 11 unique viral protein sequences and a very high concentration of positive samples involving L and NP (96.5% of the protein pairs comprising the Positive Dataset), increases the likelihood of data leakage and overfitting to the majority class. This is also evident in **Table 3**, where predictions were made based on different negative datasets. Furthermore, due to k-fold cross validation, there is considerable overlap between training and test samples, leading to data leakage and biased accuracy metrics. Instead, an independent test should be conducted to evaluate the model’s true performance. In our work, we observe a decrease in accuracy from 96% to 90% when employing independent blind testing as compared to the standard k-fold cross validation approach. Additionally, to prevent data leakage caused by similar sequences, we applied strict sequence dissimilarity using cd-hit in the training and testing datasets for Negative Datasets 2-4.

Like other DL studies [15, 16, 19, 20, 51], we initially used an unbalanced positive-to-negative dataset ratio (1:10), which resulted in an apparent increase in performance. However, this increase is misleading, as the model tends to predict the negative samples correctly when there are more negative samples, thus inflating the performance metrics. Therefore, we conducted a test using a balanced independent k-fold approach with a 1:1 ratio of positive to negative samples and observed a decrease in accuracy from 96% to 48% as compared to standard k-fold testing with a 1:10 positive to negative ratio of the dataset. We also evaluated performance using the virus-specific evaluation, treating each arenavirus protein category as a separate class and comparing the corresponding confusion matrices of all arenavirus proteins to each other. **Figure 4. and Table 5**. clearly illustrates the model’s underperformance in the minority class under standard k-fold test settings, due to factors such as overfitting and bias, compare balanced accuracy of 73.97% with independent blind testing with balanced dataset and 66.65% for standard k-fold testing.

Furthermore, the lack of interpretability in the learning process of ML model adds another layer of concern. High accuracy in such models may obscure underlying biases, indicating models capacity to recognize specific patterns within the dataset rather than its predictive power. Thus, making it hard to understand why the model makes certain predictions. This opacity can foster overconfidence in the models predictions, which is particularly risky when the model is applied to new data.

In summary, we conducted transfer learning to predict PPIs between arenaviruses and human proteins. Certainly, features derived from other virus-human PPIs provided valuable context and hypotheses that can guide further experimental investigations into arenavirus-human PPIs. However, the data scarcity of arenavirus-human PPIs presents several challenges. We demonstrated how traditional metrics like accuracy and AUPRC do not fully capture the performance of the predictive models. Our findings highlight the need for thorough evaluation methods, such as virus specific evaluation alongside independent testing, particularly for minority classes. We also suggest focusing on overall balanced accuracy and balanced accuracy for majority and minority classes instead of metrics like overall accuracy and AUPRC. This will help better assess model’s performance and identify areas for improvement. The arenavirus-human PPI dataset currently available is skewed towards certain proteins (polymerase L and nucleoprotein NP). We showed that this can cause biases towards them (majority class) and makes the model prone to overfitting. Our results emphasize the need for strategically acquiring experimental data for minority classes of understudied viruses to improve deep learning virus-human PPIs prediction.

## Acknowledgments

We would like to acknowledge Dr. Hung Do, Dr. Ken Sale and Dr. Oscar Negrete for providing scientific insights. This study was supported through funding from the Sandia National Laboratories Laboratory Directed Research and Development (LDRD) program. The authors would also like to thank the computational resources provided by the LANL Institutional Computing. This work was performed at the Los Alamos National Laboratory, which is operated by Triad National Security, LLC, for the National Nuclear Security Administration of the U.S. Department of Energy (contract 89233218CNA000001). Sandia National Laboratories is a multi-mission laboratory managed and operated by National Technology & Engineering Solutions of Sandia, LLC, a wholly owned subsidiary of Honeywell International Inc., for the U.S. Department of Energy’s National Nuclear Security Administration under contract DE-NA0003525. This paper describes objective technical results and analysis. Any subjective views or opinions that might be expressed in the paper do not necessarily represent the views of the U.S. Department of Energy or the United States Government.

