## Supplementary Material for "Challenges in predicting protein-protein interactions of understudied viruses: Arenavirus-Human interactions"

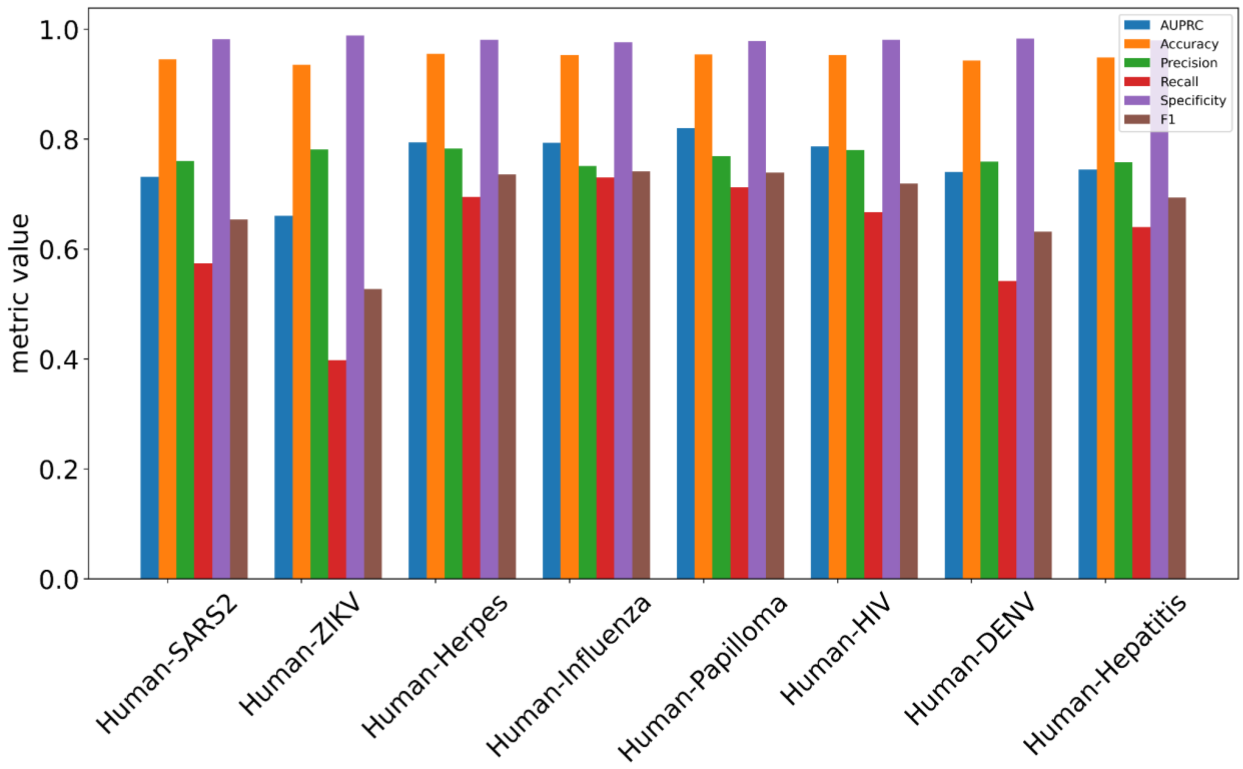


**Figure S1.** Models initially trained on various source systems [Human-DENV, Human-Hepatitis, Human-HIV, Human- SARS2, Human-ZIKV, Human-Herpes, Human-Influenza, Human-Papilloma] and finetuned on CANON dataset, performed under frozen setting.

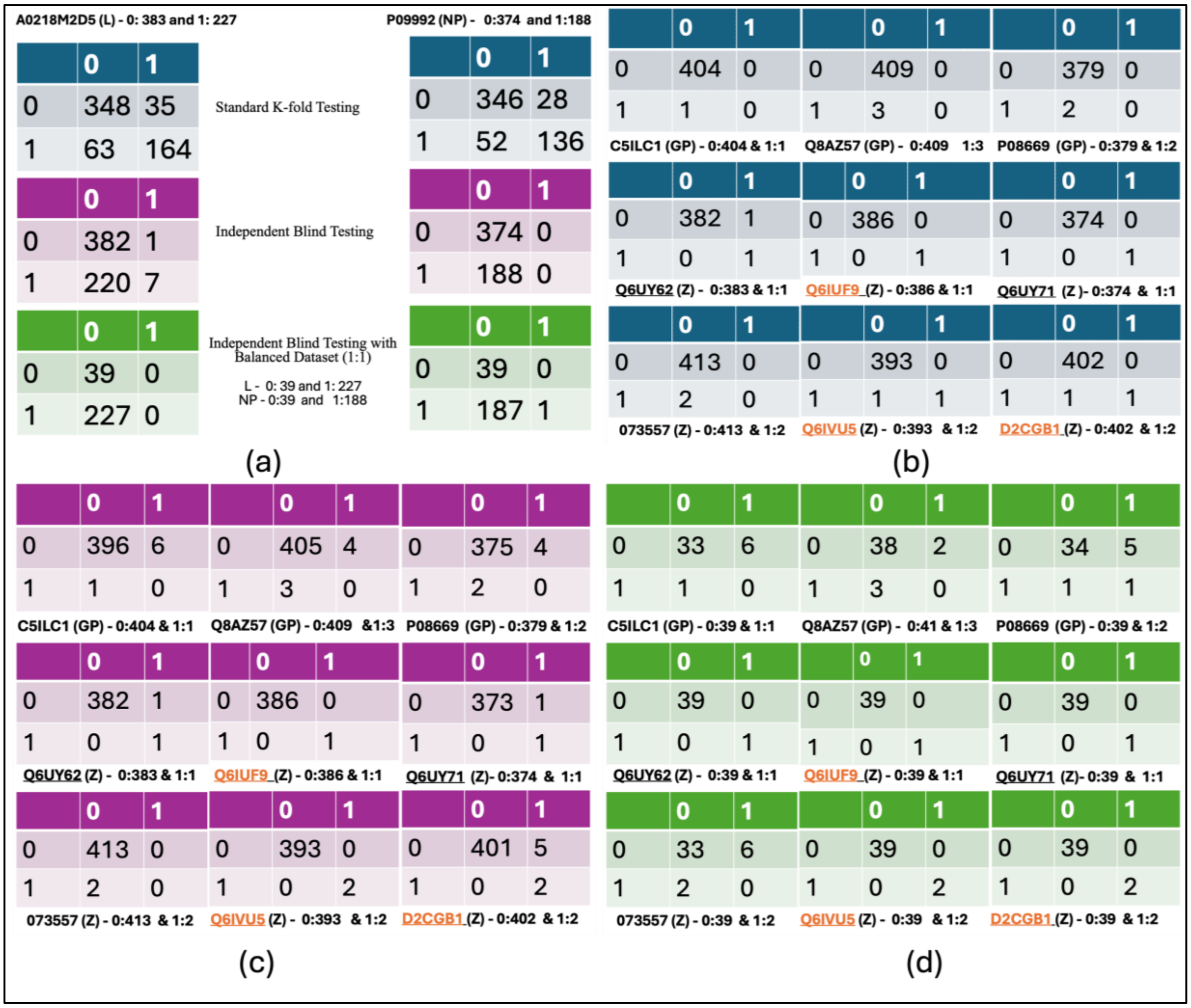


**Figure S2.** Confusion Matrices for majority and minority viral sequences from arenavirus-human PPI data (comprising the Positive Dataset) under different settings: Standard k-fold Testing, Independent Blind Testing and Independent Blind Testing with a Balanced Dataset. In each subfigure, we present the count of non-interacting samples (0: count), the count of interacting samples (1: count), and the viral protein uniprot ID. (a) presents the confusion matrix under three settings for the majority viral proteins (polymerase L and nucleoprotein NP) participating in arenavirus-human PPIs included in the Positive Dataset. (b) confusion matrix for minority viral proteins (glycoprotein GP and RING finger protein Z) under Standard k-fold Testing. (c) confusion matrix for minority viral proteins (GP and Z) under Independent Blind Testing with 1:10 negative to positive data ratio. (d) confusion matrix for minority viral sequences (GP and Z) under Independent Blind Testing with Balanced Data 1:1 negative to positive data ratio. Note: Viral proteins highlighted in orange font have 95% sequence similarity. The sequences underlined are paired with the same human protein sequence. Note, CANON dataset is used in these evaluations.

| Human Uniprot ID | Virus Uniprot ID | Arenavirus Protein | Predicted label using CANON dataset | Predicted label using RANDP dataset |
|---|---|---|---|---|
| Q8WUM4 | G3LUW8 | MOPV-NP | 1 | 1 |
| Q8WUM4 | P18140 | TCRV-NP | 1 | 1 |
| O00571 | P14239 | JUNV-NP | 1 | 0 |
| O00571 | P13699 | LASV-NP | 1 | 0 |
| O00571 | P09992 | LCMV-NP | 1 | 1 |
| Q99816 | P27588 | MARV-NP | 0 | 1 |
| Q8WUM4 | G3LUW9 | MOPV-Z | 0 | 1 |
| Q8WUM4 | Q88470 | TCRV-Z | 0 | 1 |
| Q96J02 | Q6IVU5 | JUNV-Z | 0 | 0 |
| Q96J02 | O73557 | LASV-Z | 0 | 0 |
| Q96J02 | P18541 | LCMV-Z | 0 | 0 |
| Q96J02 | C5ILC3 | Lujo-Z | 0 | 0 |
| Q96J02 | G3LUW9 | MOPV-Z | 0 | 0 |
| Q99816 | O73557 | LASV-Z | 0 | 1 |
| Q99816 | G3LUW9 | MOPV-Z | 0 | 1 |

**Supplement Table S1.** DL-mediated prediction of arenavirus-human PPIs for protein pairs that are thought to be involved in arenavirus-human PPIs based on indirect experimental associations. The pre-trained model was initially trained using HIV-Human data and then finetuned using CANON dataset and RANDP dataset. In the fourth and fifth column, 0 means predicted non-interaction, whereas 1 means predicted interaction.

| **Uniprot ID** | **Number of Negative Samples (1:10 ratio)** | **Number of Negative Samples for (1:1 ratio)** | **Number of Positive Samples** | **Protein Type** |
|---|---|---|---|---|
| A0218M2D5 | 383 | 39 | 227 | L |
| P09992 | 374 | 39 | 188 | NP |
| C5ILC1 | 404 | 39 | 1 | GP |
| P08669 | 409 | 39 | 3 | GP |
| Q8AZ57 | 379 | 40 | 2 | GP |
| Q6UY62 | 383 | 39 | 1 | Z |
| D2CGB1 | 402 | 39 | 2 | Z |
| Q6IUF9 | 386 | 39 | 1 | Z |
| Q6UY71 | 374 | 39 | 1 | Z |
| 073557 | 413 | 39 | 2 | Z |
| Q6IVU5 | 383 | 39 | 2 | Z |

**Supplement Table S2.** Total count of positive and negative samples involving each of the arenavirus protein presented in 1:10 and 1:1 positive to negative ratios.

| Arenavirus protein Uniprot ID | True Positive | False Negative | Protein Type |
|---|---|---|---|
| A0218M2D5 | 227 | 0 | L |
| P09992 | 187 | 1 | NP |
| C5ILC1 | 0 | 1 | GP |
| P08669 | 0 | 2 | GP |
| Q8AZ57 | 0 | 3 | GP |
| Q6UY62 | 1 | 0 | Z |
| D2CGB1 | 1 | 1 | Z |
| Q6IUF9 | 1 | 0 | Z |
| Q6UY71 | 1 | 0 | Z |
| 073557 | 1 | 1 | Z |
| Q6IVU5 | 1 | 1 | Z |

**Supplement Table S3.** Viral specific analysis for RANDP dataset. True positives: count of correct predictions and False Negative: count of interactions predicted incorrectly. The predictions are made by the model that was initially trained using HIV-Human data and then finetuned using RANDP dataset.
